# Done in 100 ms: Path-dependent visuomotor transformation in the human upper limb

**DOI:** 10.1101/192880

**Authors:** Chao Gu, J. Andrew Pruszynski, Paul L. Gribble, Brian D. Corneil

**Affiliations:** Department of Psychology; University of Western Ontario; London, ON, N6A 5B7; Canada; Department of Physiology & Pharmacology; University of Western Ontario; London, ON, N6A 5B7; Canada; The Brain and Mind Institute; University of Western Ontario; London, ON, N6A 5B7; Canada; The Robarts Research Institute; University of Western Ontario; London, ON, N6A 5B7; Canada

**Author notes:** Corresponding Author: Dr. Brian D. Corneil, Robarts Research Institute, University of Western Ontario, 1151 Richmond St. N, London, Ontario, Canada, N6A 5B7.

**Keywords:** Human, reaching movement, EMG, visual response, movement planning, curvature, trajectory, hand-eye coordination

## Abstract

A core assumption underlying mental chronometry is that more complex tasks increase cortical processing, prolonging reaction times. Here we show that increases in task complexity alter the magnitude, rather than the latency, of the output for a circuit that rapidly transforms visual information into motor actions. We quantified visual *stimulus-locked responses* (SLRs), which are changes in upper limb muscle recruitment that evolve at a fixed latency ∼100 ms after novel visual stimulus onset. First, we studied the underlying reference frame of the SLR, by dissociating initial eye and hand position. Despite its quick latency, we found that the SLR was expressed in a hand-centric reference frame, suggesting that the circuit mediating the SLR integrated retinotopic visual information with body configuration. Next, we studied the influence of planned movement trajectory, requiring participants to prepare and generate curved or straight reaches in the presence of obstacles to attain the same visual stimulus. We found that SLR magnitude reflected the initial planned movement trajectory, regardless of the ensuing movement curvature. Based on these results, we suggest that the circuit mediating the SLR lies in parallel to other well-studied corticospinal pathways. Although the fixed latency of the SLR precludes extensive cortical processing, inputs conveying information relating to task complexity, such as body configuration and planned movement trajectory, can pre-set nodes within the circuit underlying to the SLR to modulate its magnitude.

**SIGNIFICANCE STATEMENT:** A core assumption underlying mental chronometry is that more complex tasks increase cortical processing, prolonging reaction times. Here, we showed that increases in task complexity altered the magnitude, rather than the latency, of a circuit that rapidly transforms visual information into motor actions. We focus on *stimulus-locked responses* (SLRs), which are changes in upper limb muscle recruitment that evolve at a fixed latency ∼100 ms after novel visual stimulus onset. We showed that despite is quick latency, the circuitry mediating the SLR transformed a retinotopic visual signal into a hand-centric motor command suitable to contribute to the initial movement trajectory. We suggest that this circuit lies in parallel to other well-studied corticospinal pathways.

## INTRODUCTION

The reaction time (RT) needed to initiate a visually-guided action is a core measure in behavioral neuroscience (Luce, 1986). In humans, visually-guided reaches from a static posture typically start within ∼200-300 ms of stimulus presentation (Welford, 1980), with RTs increasing for more complex tasks that require additional cortical processing (Donders, 1969). A more precise measurement of RT can be obtained via electromyographic (EMG) recordings of limb muscle activity that circumvent the electromechanical delays that arise between the neural command to initiate a movement and movement itself (e.g. due to the arm’s inertia, Norman & Komi, 1979). In addition to the this large and well-studied volley of neuromuscular activity that initiates the movement, a brief and small burst of activity occurs time-locked ∼100 ms after novel visual stimulus presentation, regardless of the ensuing movement RT (Pruszynski *et al.*, 2010). These visual *stimulus-locked responses* (SLRs) are directionally tuned, with EMG activity increasing or decreasing for stimulus locations to which the muscle would serve as an agonist or antagonist, respectively. Furthermore, the SLR persists when movement is temporarily withheld (Wood *et al.*, 2015) or proceeds in the opposite direction (Gu *et al.*, 2016).

The SLR evolves during the earliest interval in which visual information can influence limb muscle recruitment, and its short latency limits the opportunity for extensive cortical processing. To better understand the properties of the circuit underlying this rapid sensorimotor transformation, we characterized the SLR across three different visually guided reach experiments by altering task complexity. We studied whether the SLR was expressed in an eye- or hand-centered reference frame by dissociating initial eye and hand position (Experiment 1), and the influence of different pre-planned straight or curved movement trajectories on the SLR (Experiments 2 & 3). We found that while the SLR latencies remained constant in all three experiments, changes in SLR magnitude showed that the underlying circuit rapidly transforms retinotopic visual information into a hand-centered reference frame suitable for contributing to the initial movement trajectory of the hand.

## MATERIALS AND METHODS

In total, we had 30 participants (19 males, 11 females; mean age: 26 ± 5 years SD) perform at least one of the three experiments. All were self-declared right-handed except for two left-handed males and two left-handed females. All participants had normal or corrected-to-normal vision and reported no current visual, neurological, and/or musculoskeletal disorders. Participants provided written consent, were paid for their participation, and were free to withdraw from any of the experiments at any time. All procedures were approved by the Health Science Research Ethics Board at the University of Western Ontario. Parts of the apparatus, electromyography (EMG) recording setup, and data analyses has been previously described (Wood *et al.*, 2015; Gu *et al.*, 2016).

### Apparatus and Kinematic Acquisition

Briefly, in all 3 experiments, participants performed reach movement in the horizontal plane with their right arm while grasping the handle of a robotic manipulandum (InMotion Technologies, Watertown, MA, USA). Participants sat at a desk and interacted with the robotic manipulandum with their elbow supported by a custom-built air-sled. A constant load force of 5 N to the right was applied to increase the baseline activity for the limb muscle of interest for all 3 experiments. The *x-* and *y-* position of the manipulandum was sampled at 600 Hz. All visual stimuli were presented onto a horizontal mirror, located just below the participant’s chin level, which reflected the display of a downward-facing LCD monitor with a refresh rate of 75 Hz. The precise timing of visual stimulus onsets on the LCD screen was determined by a photodiode. The mirror occluded the participant’s arm and visual feedback of the hand was given as a small red cursor.

### EMG and EOG Acquisition

EMG activities from the clavicular head of the right pectoralis major (PEC) muscle were recorded using either intramuscular (Experiment 1) and/or surface recordings (all Experiments). Intramuscular EMG activity was recorded using fine-wire (A-M Systems, Sequim, WA, USA) electrodes inserted into the PEC muscle (see Wood et al., 2015 for insertion procedure). Briefly, for each recording we inserted two monopolar electrodes ∼2.5 cm into the muscle belly of the PEC muscle, enabling recording of multiple motor units. Insertions were aimed ∼1 cm inferior to the inflection point of the participant’s clavicle, and staggered by 1 cm along the muscle’s fiber direction. All intramuscular EMG data were recording with a Myopac Junior System (Run Technologies, Mission Viejo, CA, USA). Surface EMG was recorded with doubled-differential electrodes (Delsys Inc., Natick, MA, USA), placed either near or at the same location as the intramuscular recordings. In Experiment 1, horizontal eye position was measured using bitemporal direct current electrooculography (EOG, Grass Instruments, Astro-Med Inc.). EMG and EOG data were digitized and sampled at 4 kHz.

### Data Analyses

To achieve sample-to-sample matching between kinematic and EMG data, kinematic data were up-sampled from 600 Hz to 1000 Hz with a lowpass interpolation algorithm, and then lowpass-filtered with a second-order Butterworth filter with a cutoff at 150 Hz. Off-line, EMG data was rectified, and either bin-integrated into 1 ms bins (intramuscular) or down-sampled (surface) to match the 1000 Hz sample rate. Reach reaction times (RTs) were calculated as the time from the onset of the visual stimulus (measured by a photodiode) to the initiation of the reach movement. Reach initiation was identified by first finding the peak tangential movement velocity, and then moving backwards to the closest time-point at which the velocity profile reached 8% of the peak velocity. We defined the SLR epoch as 85-125 ms after stimulus onset. Trials with RTs less than 185 ms were excluded to prevent contamination of the SLR epoch by recruitment associated with very short-latency responses (Wood *et al.*, 2015; Gu *et al.*, 2016). We also defined the voluntary movement (MOV) epoch as -20 to 20 ms around the reach RT.

To determine the normalized movement trajectory for Experiments 2 and 3, we first defined the movement duration for each trial individually. The movement duration was defined as 50 ms prior to when the hand position surpassed 2 cm from the center of the start position to 50 ms after the time when the hand position surpassed 20 cm (14 cm for the Catch Trials in Experiment 3) from the center of the start position. We then interpolated the movement duration into 101 equal time samples. Then for each normalized time sample we calculated the *x*- and *y*-position to get the normalized movement trajectory for each trial.

### SLR Detection and Latency Analysis

Based on previous works identifying the SLR (Corneil *et al.*, 2004; Pruszynski *et al.*, 2010), we used a receiver-operating characteristic (ROC) analysis to quantitatively detect the presence of a SLR. In all Experiments, we first separated the EMG activity for all correct control reaches based on visual stimulus location, and performed the following ROC analysis. For every time-sample (1 ms bin) between 100 ms before to 300 ms after visual stimulus onset, we calculated the area under the ROC curve. This metric indicates the probability that an ideal observer could discriminate the side of the stimulus location based solely on EMG activity. A value of 0.5 indicates chance discrimination, whereas a value of 1 or 0 indicates perfectly correct or incorrect discrimination, respectively. We set the thresholds for discrimination at 0.6; these criteria exceed the 95% confidence intervals of data randomly shuffled with a bootstrap procedure (Chapman & Corneil, 2011). The earliest discrimination time was defined as the time after stimulus onset at which the ROC was above 0.6 and remained above that threshold for at least 5 out of the next 10 samples. Based on the ROC analyses we defined the SLR epoch as from 85 to 125 ms after visual stimulus onset and categorized any participant with a discrimination time <125 ms as having a SLR (SLR+ participant). Across the 5 experiments we could reliably detect a SLR in 24 out of 30 participants (∼80% detection rates). This rate is comparable to previous reports of the SLR detection on the limb with either intramuscular and surface recordings in this setup (Wood *et al.*, 2015; Gu *et al.*, 2016). To determine the onset latency of the SLR on the upper limb, we used the same procedure as previously described for SLR on neck muscle activity (Goonetilleke *et al.*, 2015). Briefly, we used the same time-series ROC mentioned above and fit a two-piece piecewise linear regression (Cashaback *et al.*, 2013). The first linear regression is based on baseline activity preceding any SLR (from 0 to 80 ms after stimulus onset) and the second linear regression is based on activity for candidate inflection point to the peak of the SLR (max ROC value in an interval from 80 to 140 ms). The inflection point was determined as the latency that minimized the sum of the squared error between the observed ROC curve and the two linear regressions. Relative to the ROC value at the inflection point, the onset latency was the time where the ROC increased by 0.05 for the next 5 out of 10 samples.

### Experiment 1: Reference Frame Task

To initiate each trial, participants (*N* = 7/8; 7 SLR+ participants) brought the cursor into a starting hand position (**Fig. 1a**, green circle). After a randomized (0.5 - 1 sec) delay, participants had to look towards the starting eye position (red circle). 3 different initial positions were possible: either the hand and the eye were in line with the participant’s midline (Position 1), or the hand was 10 cm to the right and the eye was 10 cm to left of midline (Position 2), or vice versa where the hand was 10 cm to the left and the eye was 10 cm to the right of midline (Position 3). After another randomized (1 - 1.5 sec) delay, a black visual stimulus appeared concurrently with the offset of both the starting hand and eye position stimuli. This served as the go cue to make a coordinated hand-eye movement towards the black visual stimulus. The black stimulus could be in 1 of 3 possible locations: either at the midline (Stim_C_) or 20 cm to the left (Stim_L_) or right of midline (Stim_R_). If the participant moved their hand outside of the starting hand position at any point prior to the onset of the black stimulus, the trial was aborted and reset. 1 second after the onset of the black stimulus, the next trial started. Participants performed 64 trials for each of 9 different conditions across 8 different blocks. For Stim_C_ in Position 1 participants were not required to move. To analyze the data during the presumed MOV epoch on such trials, we assumed that the RT for these trials would be from a similar distribution of RT as Stim_L_ and Stim_R_ reach movements. Thus, we randomly assigned a reach RT for each Stim_C_ trial from the pooled RT of Stim_L_ and Stim_R_ in Position 1.

**Figure 1:**
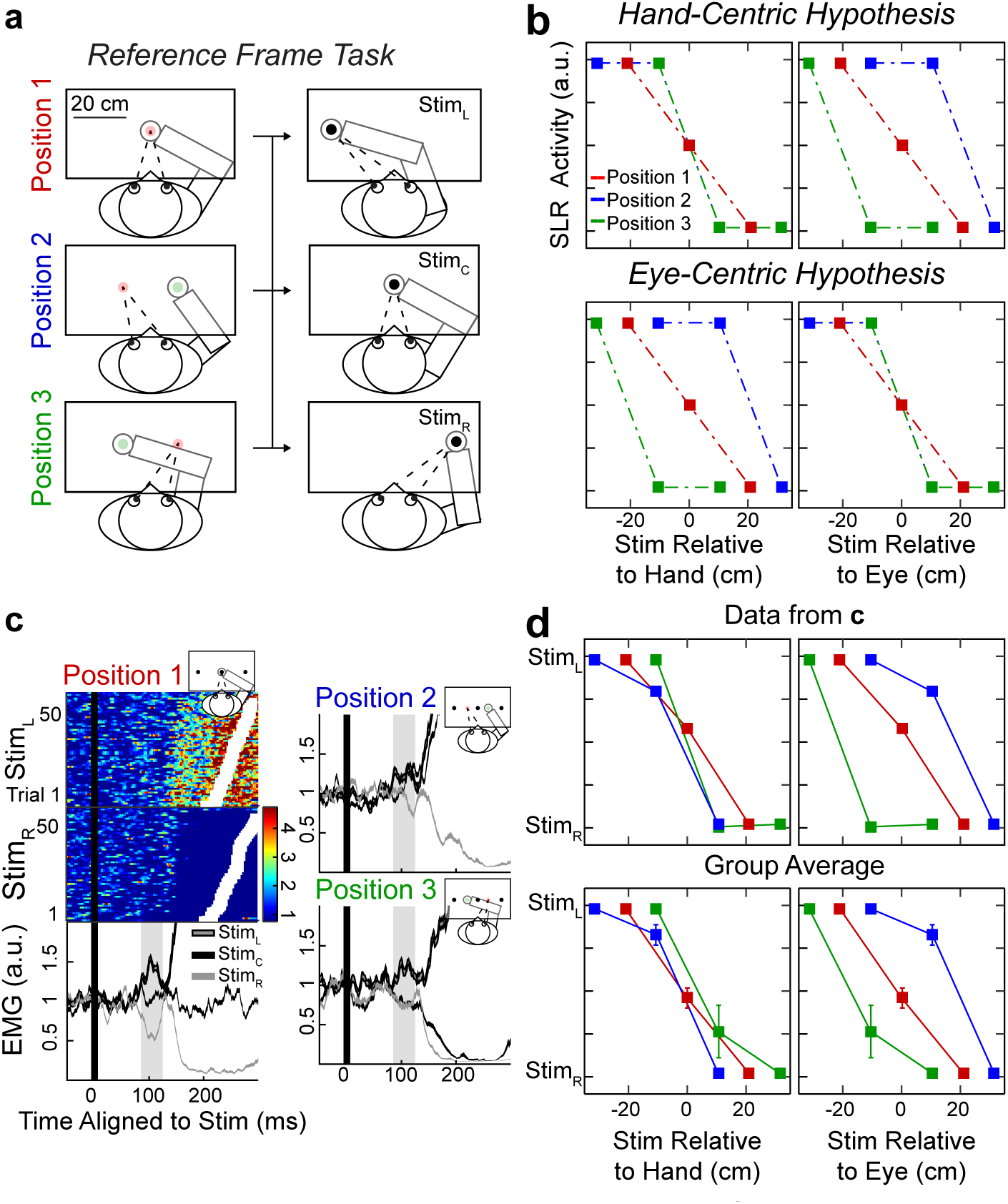
The SLR generates a motor command toward the visual stimulus in a hand-centric reference frame. **a.** Experimental paradigm. Participants started in 1 of 3 different initial positions, and were instructed to move their eyes and right hand to a black visual stimulus. **b.** These various initial positions and stimulus locations allowed us to predict SLR magnitude as either a function of stimulus location relative to either the hand (*hand-centric*, top) or the eye (*eye-centric* reference frame, bottom panels). **c.** Individual and mean EMG activity from a participant. The color subpanels are individual Stim_L_ and Stim_R_ trials from Position 1. Each row represents EMG activity from a single trial, with all trials aligned to stimulus onset (black line) and sorted based on reach RT (white squares). All other subpanels represent mean ± SEM EMG activities for correct trials, segregated by initial position and stimulus location. Shaded boxes indicate the SLR epoch. **d.** The participant for **c.** (top) and the group (*n =* 7, bottom panels) mean adjusted SLR magnitudes conform to the prediction of a *hand-centric* reference frame.

### Experiment 2: Obstacle Task

Each trial began with the appearance of a start position stimulus; on 2/3^rd^ of all trials the gray visual obstacle was presented concurrently. No obstacle was presented on the other 1/3^rd^ of trials, which served as a control condition. Two different sets of obstacles could appear, either a horizontal bar or two upside-down L-shape obstacles (**Fig. 2**). To initiate the trials participants (*N* = 15/20 SLR+) moved the cursor into the start position. After a variable delay (1 - 1.25 sec) a black peripheral stimulus appeared 20 cm from the position, at either a left-outward (135° CCW from straight right) or right-outward (45° CCW) location. The start position was extinguished simultaneously with the presentation of the peripheral stimulus. Participants then had to move the cursor as quickly as possible to the peripheral stimulus. Each participant performed 4 blocks; in 2 blocks participants were instructed to avoid the gray obstacles while in the other 2 blocks they were instructed to reach through the obstacles when reaching for the peripheral stimulus. The order of instruction was counterbalanced across our participants. Each block consisted of 150 trials in total, with 25 trials for each of the 6 different conditions.

**Figure 2:**
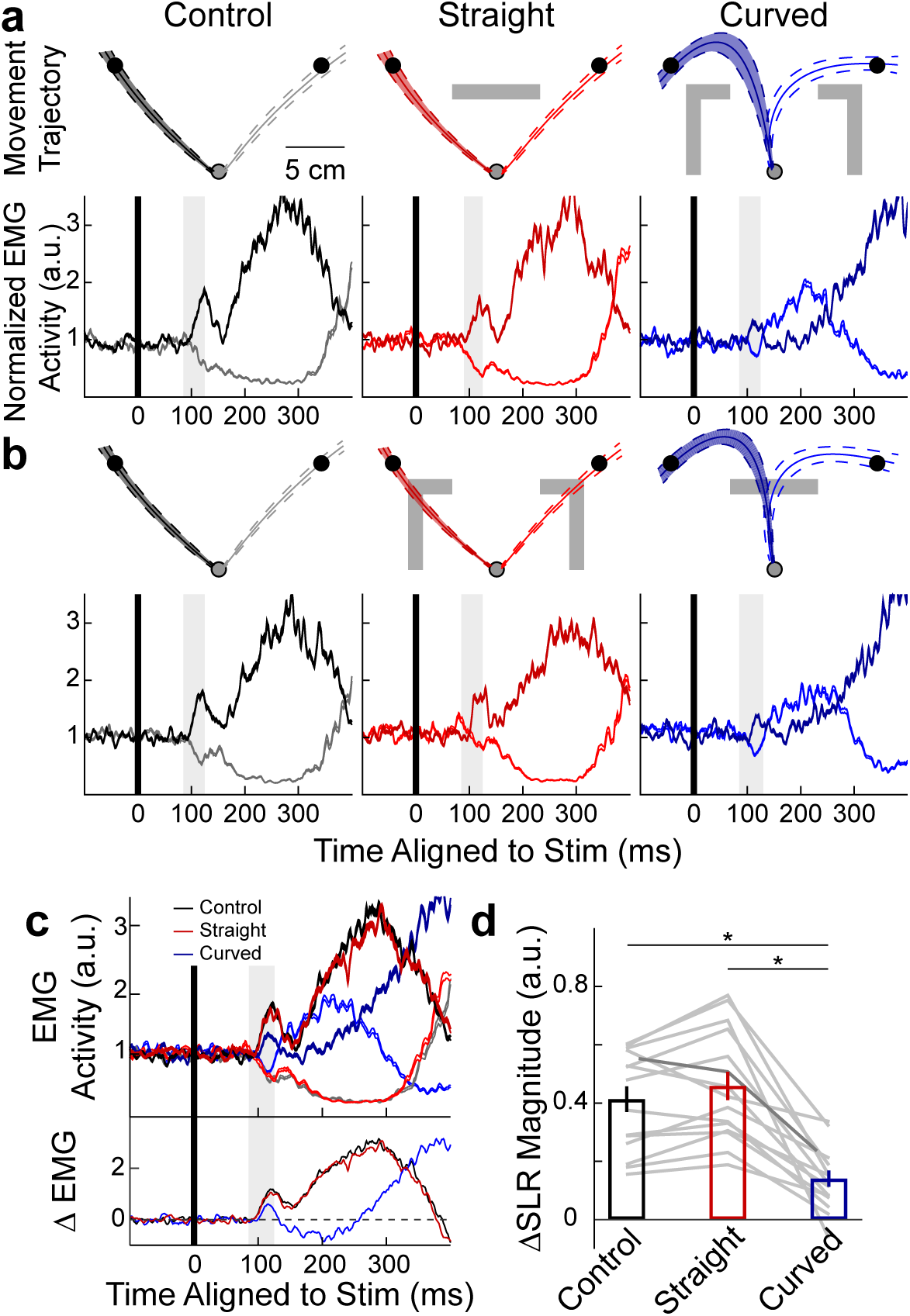
Decreased SLR magnitude for curved compared to straight reaches to the same visual stimulus. **a, b.** Kinematic and EMG data from a participant during the obstacle task when instructed to avoid or reach though the visual obstacle (gray rectangles). Left-outward (dark) and right-outward (light shaded contours) reach trials were segregated by movement trajectory: reaches with no obstacles (Control– black), straight reaches with obstacles (Straight – red), or curved reaches with obstacles (Curved – blue). Top panels show the mean ± SD normalized movement trajectories for each condition, while the bottom panels show the corresponding mean ± SEM EMG activities aligned to stimulus onset, with the SLR epoch highlighted (shaded boxes). **c.** Top subpanel shows the same EMG data as **a** and **b.**, but combining EMG data for the 3 difference movement trajectories regardless of visual obstacle and task instruction. Bottom subpanel shows the difference in mean EMG activity (ΔEMG) between left-outward and right-outward reach trials for the 3 trajectories. **d.** Group mean ± SEM of the ΔEMG during the SLR epoch (ΔSLR magnitude) for the 3 different movement trajectories. Each gray line represents an individual participant, with the darker line representing data from the participant in **c**. * *P* < 0.0001.

### Experiment 3: Choice Task

Each trial began with the appearance of a start position stimulus and a gray obstacle (**Fig. 3a**). To initiate the trials participants (*N* = 14/15 SLR+) moved the cursor into the start position. After a variable delay (1 - 1.25 sec) the start position was extinguished simultaneously with the presentation of the peripheral black visual stimulus. On Test Trials (2/3^rd^ of all trials) the peripheral stimulus was presented 20 cm left-outward from the start position (135° CCW), while in Catch Trials (1/3^rd^ of all trials) the peripheral stimulus was presented 14 cm from the start position directly outward (90° CCW) or leftward (180° CCW) with equal likelihood. Participants were instructed to move the cursor as quickly as possible to the peripheral stimulus, while avoiding the gray obstacle by choosing the shortest movement trajectory. The shape of the gray obstacle varied on a trial-by-trial basis but the overall area remained constant. The obstacle shape displayed was based on an adaptive estimation of the psychometric function for each participant. We assumed that the psychometric function of the choice of the movement trajectory around the obstacle took the form of a logistic function (**Eq. 1**):

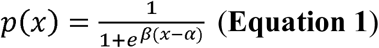

in which *x* is the shape of the obstacle (ranging from a purely horizontal bar, *x* = -68, through L-shaped obstacles, through to a vertical bar, *x* = 68; for shapes see x-axis of **Fig. 3b**); *p(x)* indicates the probability of leftward curved reach around the obstacle for the given midpoint of the obstacle; and *α* and *β* are the threshold and the slope of the logistic function. To estimate this function, we used a modified updated maximum likelihood procedure (Shen & Richards, 2012), with the parameter space consisting of a grid of *α* and *β* values. The *α* parameter spanned 69 values ranging from -68 to 68 in 2 unit increments. The *β* parameter spanned value ranging from 0 to 0.5 in 0.05 increments. A uniformed prior (*α*, *β* = 0) was used for the 1^st^ block, while subsequent blocks used the estimated parameters from the last trial of the previous block. To initialize each block, the first 5 trials had obstacles that were at the: 0^th^, 100^th^, 50^th^, 25th, and 75^th^ percentiles (x = -68, 68, 0, -34, 34 unit, respectively). Afterwards, the obstacle shape was set either at the estimated threshold, *p(x)* = 0.5, or at either the lower, *p(x)* = 0.25, or upper deflection, *p(x)* = 0.75, deflections in a pseudorandom order at a 2:1:1 ratio. Test, Catch leftward, and Catch rightward Trials were also presented in a pseudorandom order at a 4:1:1 ratio, respectively. Each participant performed 6 blocks, except for 1 whom performed 5 blocks, with each block consisting of 197 trials: 5 initial trials, 128 Test, 32 Catch leftward and rightward Trials. All participants had at least 100 correct Test Trials for the threshold visual obstacle, at which *p*_*(leftward)*_ was closest to 0.5.

**Figure 3:**
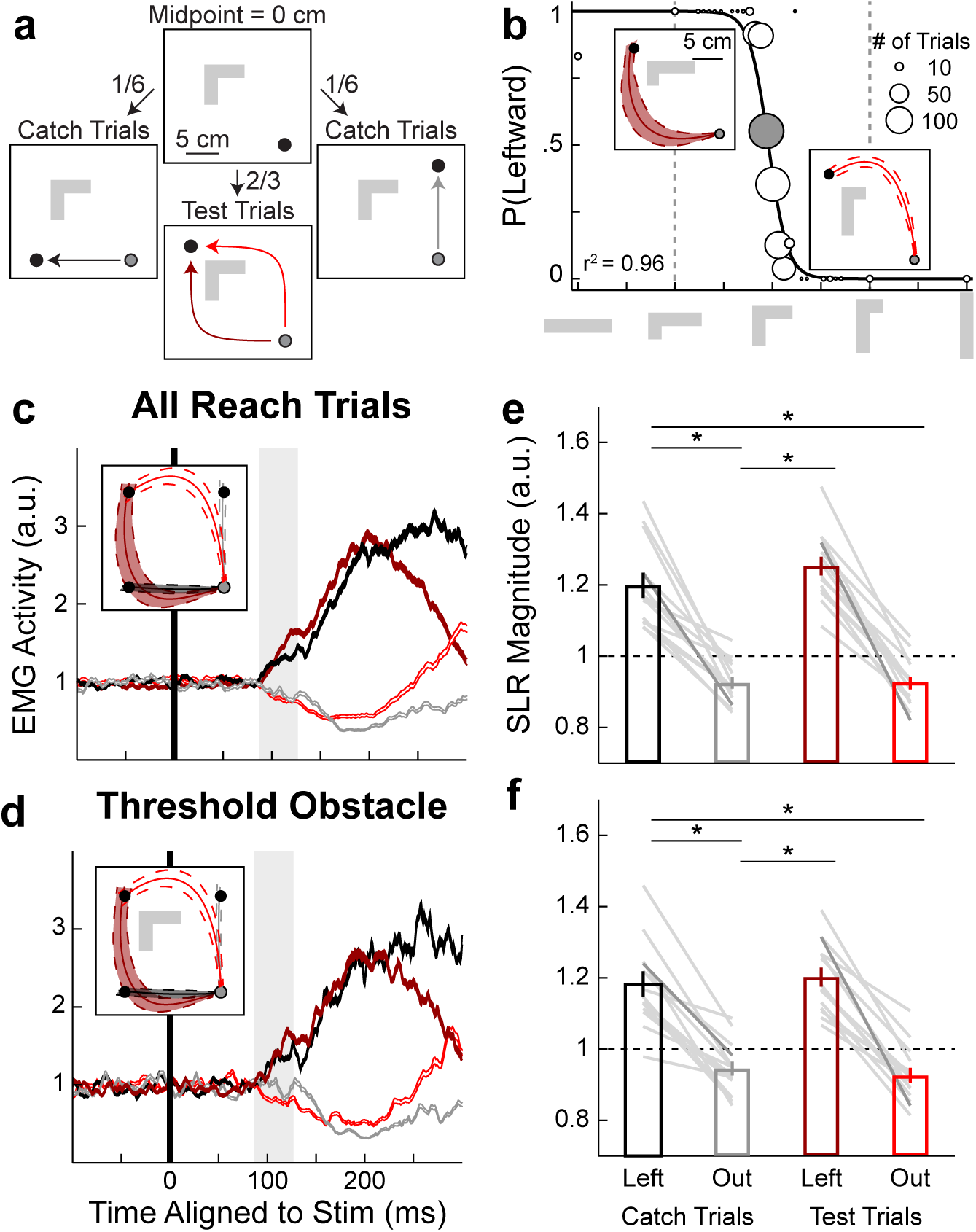
SLR magnitude modulated by pre-planned movement trajectory. **a.** Experiment paradigm. Participants were instructed to reach to a visual stimulus using the shortest movement trajectory while avoiding an obstacle. The shape of the obstacle varied on a trial-by-trial basis (see **METHODS** for exact details). For Test Trials, participants made either initially curved leftward (dark red) or outward (light red) reaches toward a left-outward visual stimulus. For Catch Trials, participants reached straight leftward (black) or outward (gray). **b.** Behavioral performance for all Test Trials from a participant. The probability of an initial leftward curved reach is plotted as a function of obstacle shape. Insert shows the mean + SD normalized movement trajectories for two different obstacle shapes. Black line is the best fit of a logistic function (**Eq. 1**) to the participant’s behavior. **c, d.** The participant’s mean EMG + SEM different reach types for all reach trials (**c**) and for the threshold obstacle (**d**, shaded circle from **b**) aligned to stimulus onset. **e, f.** Group mean + SEM of SLR magnitudes for all reach trials (**e**) and for the threshold obstacle (**f**) for the 4 different reach conditions. Each gray line represents an individual participant, with the darker line indicating data from the participant in **b**. * *P* < 10^-6^.

### Experimental Design and Statistical Analysis

All statistical analyses were performed with custom-written script in Matlab (version R2014b, Mathworks Inc, Natick, MA, USA). In Experiment 1, the within subject analysis was a 2-way ANOVA, with the mean factors of start position and stimulus location, while the between subject analysis was a 1-way ANOVA for the mean adjusted EMG activity of each start position. In Experiment 2, the within subject analysis was a 2-way ANOVA, with the mean factors of stimulus location and movement trajectory. while the between subject analysis was a 1-way ANOVA of the normalized EMG activity for movement trajectory. Finally, in Experiment 3, for both within and between subject analyses, we performed a 2-way ANOVA, with the mean factors of reach direction and movement trajectory. The level of significance was set to *P* < 0.05 at the group level, and *P* < 0.05 post-hoc test Tukey’s HSD corrected.

## RESULTS

In total 30 participants took part in at least one of the three experiments (42 separate sessions in total). In all experiments, participants used their right hand to interact with the handle of a robotic manipulandum. We measured EMG activity from the right PEC muscle while participants performed horizontal planar reaching movements. A reliable SLR was detected in 24 out of 30 participants (SLR+, ∼80%) participants (see **Methods** for detection criteria). This SLR detection rate of ∼80% was similar to our previous studies (Wood *et al.*, 2015; Gu *et al.*, 2016). Data from participants that did not exhibit a SLR were excluded from all subsequent analyses.

### The SLR encodes stimulus location relative to hand, not eye, position

Although previous studies have reported that SLRs are tuned to the position of the visual stimulus (Pruszynski *et al.*, 2010; Wood *et al.*, 2015; Gu *et al.*, 2016), these studies did not manipulate the initial position of the eyes and hand and thus could not differentiate whether the SLR encoded stimulus position relative to the eye or hand. The underlying reference frame of the SLR may start to reveal the underlying neural circuitry since many fMRI and neurophysiological studies have shown that visual stimuli can be encoded in different reference frames throughout the parietal and motor cortices (Batista *et al.*, 1999; Buneo *et al.*, 2002; Medendorp *et al.*, 2003; Crawford *et al.*, 2004; Pesaran *et al.*, 2006).

In Experiment 1, we assessed if the SLR encoded stimulus location relative to the eye (an *eye-centric*) or the hand position (a *hand-centric* reference frame). Participants (*N* = 7/8, 7 SLR+ participants) began each trial in 1 of 3 initial positions (**Fig. 1a**), with either the hand and eye in line with the participant’s midline (Position 1, red), with either the hand 10 cm right and the eye 10 cm left of midline (Position 2, blue), or with the hand 10 cm left and the eye 10 cm right of midline (Position 3, green). Participants then made a coordinated hand-eye movement towards a black visual stimulus that appeared either 20 cm left (Stim_L_), 20 cm right (Stim_R_), or at the midline (Stim_C_). These various initial positions and stimulus locations allowed us to predict SLR magnitude as a function of stimulus location relative to either the hand or eye position (**Fig. 1b**). Note that if the SLR is aligned in the corrected reference frame, we predict an overlapping SLR for all three different starting positions.

**Figure 1c** shows a participant’s EMG activity aligned to visual stimulus onset (black line) from all 3 initial positions. Trials were segregated based on initial position and visual stimulus location. EMG activity was normalized to background activity (mean EMG activity 41 ms prior to stimulus onset) for each position separately. In Position 1, similar to previous reports, we observed a reliable difference in SLR magnitude (shaded box spanning 85-125 ms after stimulus onset, left bottom panel) between Stim_L_ and Stim_R_ trials (2-way ANOVA – start position and stimulus location, interaction effect, *F*_(4,553)_ = 4.88, *P* = 0.0007, post-hoc Tukey’s HSD, *P* < 10^-7^). This increase and decrease in EMG activity can also be seen on individual EMG traces from the Stim_L_ and Stim_R_ trials, respectively (top-left and middle-left panels). The SLR was relatively brief and evolved before the much larger change in EMG activity associated with either the leftward or rightward reach movement (RTs denoted by white squares). The SLR persisted in the other 2 initial Positions, with SLR magnitude being reliably greater for Stim_L_ compared to Stim_R_ trials (**Fig. 1c**, *P* = 0.0002 and *P* < 10^-6^, for Positions 2 and 3, respectively). Across our participants, we found no difference in the onset latency of the SLR for Stim_L_ and Stim_R_ trials between when the hand and eye started in the same (Position 1, mean ± SEM latency = 87.4 ± 1.2 ms) versus different locations (Positions 2 and 3, 86.7 ± 2.2 ms; paired t-test, *t*_6_ = 0.28, *P =* 0.79), even though the median RTs were slightly shorter when the eye and hand started at the same (272.4 ± 9.4 ms) versus different positions (286.0 ± 11.2 ms; paired t-test, *t*_6_ = –3.1, *P* = 0.02).

In Positions 2 and 3, the Stim_C_ trials can be used to differentiate between hand-centric and eye-centric reference frames, since the stimulus falls between the initial positions of the hand and eye. In Position 2, SLR magnitude increased by an equal amount for both Stim_C_ and Stim_L_ trials (*P* = 0.89), when the stimulus fell to the left of the hand. In Position 3, SLR magnitude decreased by an equal amount for both the Stim_C_ and Stim_R_ trials (*P* = 0.99), when the stimulus fell to the right of the hand. Thus, for this participant, the pattern of SLR responses is consistent with a hand-centric reference frame. To account for the differences in SLR magnitude for each Position and across our participants, we scaled the SLR magnitude for Stim_C_ trials based on the SLR magnitudes observed for Stim_L_ and Stim_R_ trials (+1, -1 a.u., respectively). This allowed us to test our data against the 2 initial predictions, expressing the adjusted SLR magnitudes aligned to stimulus location relative to either the hand or eye position for this participant (top row, **Fig.1d**) and across the group (bottom row). Our results clearly indicate that the SLR is encoded in a hand-centric reference frame (compare to the *hand-centric hypothesis* in **Fig. 1b**). Across the group, we found reliably greater SLR magnitudes for Stim_C_ trials in Position 2 compared to Position 3 (1-way ANOVA - position, *F*_(2,18)_ = 7.64, *P* = 0.004, post-hoc Tukey’s HSD, *P* = 0.003). We found a similar response pattern during the MOV epoch, where Position 2 evoked a greater MOV response compared to Position 3 (*F*_(2,18)_ = 302.8, *P* < 10^-13^, post-hoc Tukey’s HSD, *P* < 10^-8^). This result suggests that, despite its short latency, the circuit mediating the SLR rapidly integrate visual stimulus location and the underlying body position, generating a motor command in a hand-centric reference frame.

### Movement trajectory influences SLR magnitude for reaches to the same visual stimulus

Given that the SLR encoded the visual stimulus relative to the current hand position, we next examined if the SLR simply encoded visual stimulus location in space, or if it is influenced by the planned movement trajectory. To start differentiating these two possibilities, in Experiment 2, participants (*N* = 15/20 SLR+) performed either curved or straight reaches to two potential visual stimulus locations. In different blocks, participants were instructed to either avoid or reach through different visual obstacles to attain the left-outward or right-outward visual stimulus. Except for control trials without the obstacle, obstacles were present at trial onset so that participants could plan their trajectory to the two potential stimulus locations. **Figure 2** shows the mean normalized movement trajectories and EMG activities when a participant either avoided (**Fig. 2a**) or reached through (**Fig. 2b**) the obstacle. Trials were categorized based on movement trajectories: straight with no obstacle (Control - black), straight either avoiding or reaching through an obstacle (Straight - red), or curved either avoiding or reaching through an obstacle (Curved - blue). When categorized this way, we found no reliable difference in mean SLR magnitude across our sample for avoiding compared to reaching through the different visual obstacles (3-way ANOVA - stimulus location, movement trajectory, and instruction, main effect for instruction *F*_(1,175)_ = 0.03, *P* = 0.85). Thus, all subsequent analyses examined mean SLR magnitudes as a function of stimulus location and movement trajectory.

**Figure 2c** shows the same participant’s EMG data, but now with the EMG activity combined between the two different instructions (top panel). To compare the difference in SLR magnitude (ΔSLR magnitude) between curved and straight reach trials, we calculated the mean EMG difference between left-outward and right-outward stimulus locations (bottom panel) during the SLR epoch for the 3 different movement trajectories. Once again, across our participants we could not find a difference in SLR latency between Straight and Curved trajectories (95.1 ± 1.6 ms and 99.8 ± 2.8 ms, paired t-test, *t*_14_ = –1.9, *P* = 0.07). Note the increase in SLR latency compared to Experiment 1 is probably due to the change in stimulus locations, as left- and right-outward are not the PD and non-PD of the SLR (Pruszynski *et al.*, 2010). Instead, we did find a reliable decrease in ΔSLR magnitude for Curved reaches compared to both Control and Straight reaches (**Fig. 2d**, 1-way ANOVA - movement trajectory, *F*_(2,42)_ = 17.53, *P* < 10^-5^, post-hoc Tukey’s HSD, *P* = 0.001 and *P* < 10^-4^, respectively). The decrease in ΔSLR magnitude between Curved and Straight reaches was likely not due to a potential confound of increased RTs (Pruszynski *et al.*, 2010; Gu *et al.*, 2016), as Curved reaches had shorter median RTs than Straight reaches (268.1 ± 6.6 ms and 277.3 ± 6.4 ms, respectively, paired t-test, *t*_14_ = 2.76, *P* = 0.015). Next, we re-examined the EMG activity during the MOV epoch. As expected given the initial outward trajectory for the Curved reaches, which is associated with less PEC muscle recruitment, EMG activity for the MOV response was also attenuated for Curved compared to Control and Straight reaches (*F*_(2,42)_ = 10.1, *P* = 0.0003, post-hoc Tukey’s HSD, both *P* = 0.001). This result suggests that the SLR is not simply encoding the spatial location of a stimulus, but rather that the SLR is also influenced by the movement trajectory that is being planned to attain that stimulus.

### Initial movement trajectory, not task demands, influences SLR magnitude for curved reaches

A potential confound in Experiment 2 was the overall difference in task demand related to planning a curved versus a straight reach movement. Previous work has shown that curved reaches were more task demanding than straight point-to-point reaches (Wong *et al.*, 2016), and we previously shown that SLR magnitude decreased during a more demanding task (Gu *et al.*, 2016). In Experiment 3, we controlled for task demand by having participants (*N* = 14/15 SLR+) perform two different curved reach trajectories to attain the same visual stimulus (**Fig. 3a**). At the beginning of each trial a visual obstacle, which participants were instructed to avoid, was shown. In Test Trials, participants made either an initially leftward (dark) or outward (light red) curved movement to a left-outward stimulus. We varied the shape of the obstacle on a trial-by-trial basis (see **METHODS:** *Experiment 3* for exact detail). **Figure 3b** shows the probability of a leftward Curved reach as a function of the possible obstacle shape. The obstacle where p_(leftward)_ ≈ 0.5 was preferentially sampled was termed the threshold obstacle (filled circle). In addition, we interleaved Catch Trials so that participants made straight leftward (black) and outward (gray) movements that had similar initial trajectories as the curved movements (see insert for movement trajectories in **Fig. 3c, d**). Once again, we found no difference in the SLR latency for Curved vs Catch trials (95.9 ± 1.3 ms and 101.3 ± 4.1 ms, respectively, paired t-test, *t*_13_ = 1.33, *P* = 0.21).

To analyze this dataset, we first pooled all correct trials together regardless of the obstacle’s shape for a single participant. On Catch Trials, the SLR magnitude was greater for leftward compared to outward straight reaches (**Fig. 3c**, 2-way ANOVA – initial direction and trajectory type, interaction effect, *F*_(1,1113)_ = 5.31, *P* = 0.02, post-hoc Tukey’s HSD, *P* < 10^-8^). Similarly, on Test Trials the SLR magnitude was also greater for leftward compared to outward curved reaches (*P* < 10^-8^). When we compared reaches with the same initial movement trajectory (straight vs curved reaches), we found no reliable difference in SLR magnitudes for both initially leftward and outward reaches (*P =* 0.15 and *P* = 0.68, respectively). To further examine the influence of the planned movement trajectories on the SLR magnitude, we next examined trials at the threshold obstacle, where the exact same visual obstacle was presented and the participant generated leftward or outward curved movement trajectories approximately half the time (p_(leftward)_ = 0.55, filled circle in **Fig. 3b**). As before, the SLR magnitude was greater for leftward versus outward reaches for both Catch and Test Trials, (**Fig. 3e**, 2-way ANOVA, interaction effect, *F*_(1,279)_ = 41.4, *P* < 10^-9^, post-hoc Tukey’s HSD, *P* < 10^-8^ and *P* = 0.03, respectively). Furthermore, the SLR magnitudes were not different for straight versus curved reaches with the same initial trajectory (*P* = 0.31 and *P* = 0.78, for initially leftward and outward reaches, respectively).

We observed the same pattern of SLR magnitude modulation based on initial movement trajectory across our participants: SLR magnitude was greater for leftward versus outward reaches when pooled for all obstacles (**Fig. 3e**) and for the threshold obstacle (**Fig. 3f**, 2-way ANOVA, main effect of direction, *F*_(1,52)_ = 160.44 and 104.64, both *P* < 10^-13^, post-hoc Tukey’s HSD, all *P* < 10^-9^, respectively). Again, we found no differences in SLR magnitude for the same initial movement trajectory (all *P* > 0.38). Thus, even when we controlled for task demand by having participants perform curved reaches with different initial trajectories to the same visual stimulus location, we found that the SLR was still modulated by the initial movement trajectory. Likewise, when we re-examined the data for the MOV response we found increased PEC muscle recruitment for leftward versus outward movement trajectories (2-way ANOVA, main effect of direction, *F*_(1,52)_ = 129.38 and 138.43, both *P* < 10^-14^, post-hoc Tukey’s HSD, all *P* < 10^-9^, respectively). Thus, SLR magnitude for the same visual stimulus is modulated by the initial planned movement trajectory.

### SLR magnitude during Catch Trials were modulated based on the pre-planned movement

Finally, to further demonstrate that the SLR magnitude was modulated based on the pre-planned movement we further examined the SLR on Catch Trials. Recall that Catch trials were randomly interleaved throughout the experiment, appearing at the Leftward or Outward locations regardless of obstacle shape. Given that the obstacle was present from the start of the trial, Catch trials could be classified as being either congruent (i.e. the pre-planned movement was in the same direction as the Catch Trial) or incongruent (i.e. in the opposite direction; **Fig. 4a**). For example, obstacles more horizontal than the threshold obstacle (light grey shaded region in **Fig. 4a**) were congruent for Leftward and incongruent for Outward Catch Trials. In contrast, obstacles more vertical than the threshold obstacle (non-shaded region in **Fig. 4a**) were congruent to Outward and incongruent to Leftward Catch Trials.

**Figure 4:**
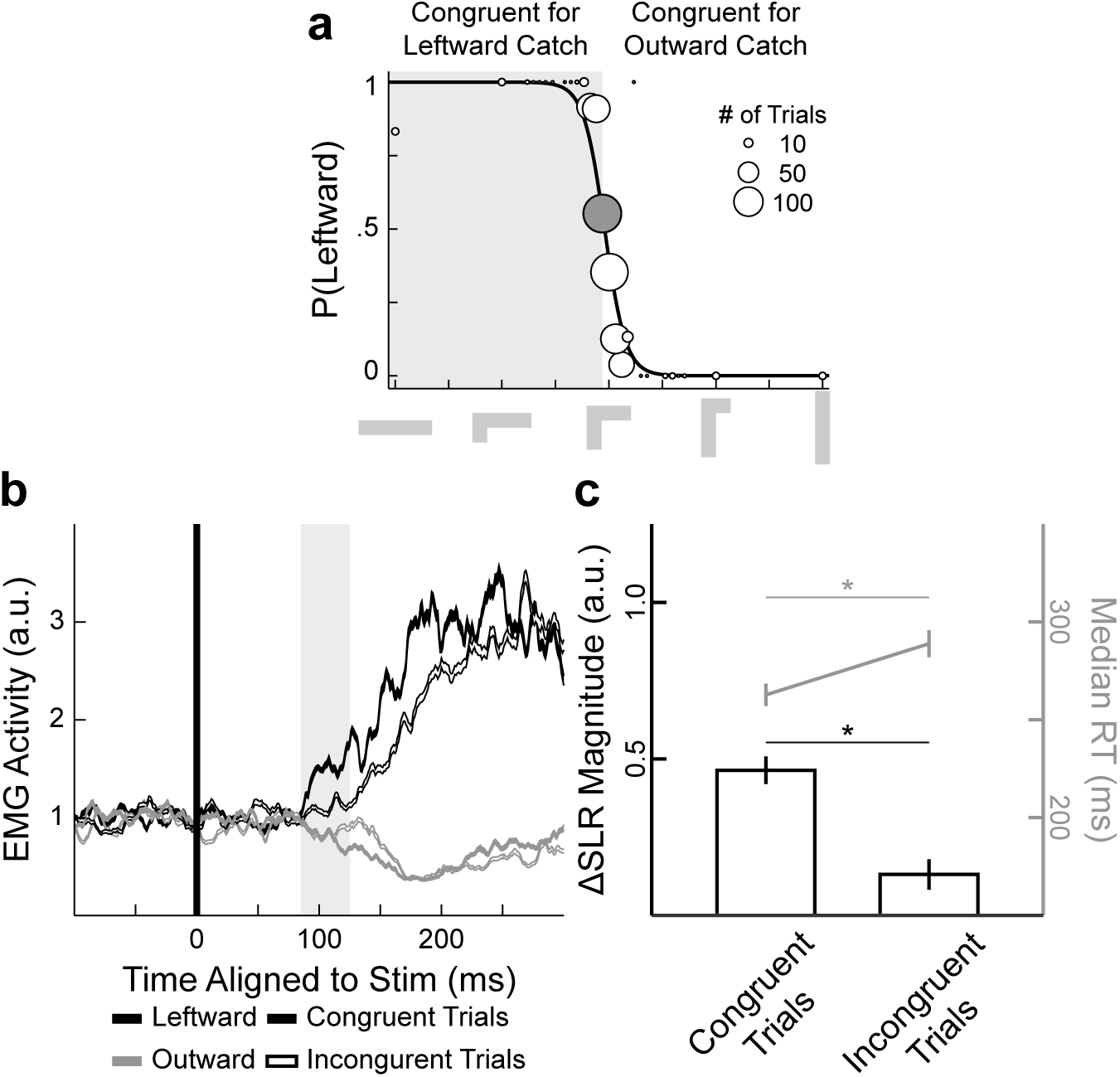
SLR magnitude and RT modulation for Catch Trials based on obstacle shape. **a.** Catch Trials were separated into Congruent and Incongruent Catch Trials. Leftward Congruent and Incongruent Trials were any trials with an obstacle more horizontal (gray shaded region) and vertical (non-shaded region) than the threshold obstacle (filled circle), respectively. For Outward Trials that was the opposite, where Congruent and Incongruent Trials were more vertical and horizontal than the threshold obstacle. **b.** Mean ± SEM for Congruent (filled) and Incongruent (open) Catch Trials sorted by either Leftward (black) or Outward (gray) stimulus location from the same participant as in **Fig. 3.c.** The mean ± SEM ΔSLR magnitude (Leftward – outward, black) and median RT (gray) for both Congruent and Incongruent Trials across our participants. * *P* < 10^-4^.

**Figure 4b** shows the mean EMG activity for all Catch Trials when we separated for both direction (Leftward – black and Outward – gray) and congruency (Congruent – filled and Incongruent – open). Note that we observed a reliable difference in EMG activity during the SLR epoch for both Congruent and Incongruent Trials, but the magnitude of the SLR was smaller for incongruent trials. **Figure 4c** shows the mean ΔSLR magnitude (black bars, Leftward – Outward Catch Trials) and median RT (gray line) across all our participants, for Congruent and Incongruent trials. We found a reliably larger ΔSLR magnitude for Congruent compared to Incongruent Trials (paired t-test, *t*_13_ = 6.88, *P* < 10^-4^), but the Incongruent ΔSLR magnitude was still present (1-sample t-test, *t*_13_ = 2.71, *P* = 0.018). Consistent with the changes in ΔSLR magnitude, we also observed difference in the ensuing RT, where participants had substantially shorter RTs for Congruent compared to Incongruent Trials (262.8 ± 5.7 and 288.9 ± 6.9 ms, respectively, paired t-test, *t*_13_ = -6.55, *P* < 10^-4^).

## DISCUSSION

Here, we characterized the visual stimulus-locked response (SLR) on the pectoralis major muscle during three different visually-guided reach tasks. Previous work has shown that the SLR is the first wave of muscle recruitment that is evoked by the onset of a novel visual stimulus, occurring within 100 ms of stimulus onset and preceding the larger volley of EMG activity associated with movement initiation (Pruszynski *et al.*, 2010; Wood *et al.*, 2015). The design of each task was based on earlier work conducted in either human or non-human primates, allowing for a direct comparison of SLR measurements to neurophysiological and behavioral concepts of sensorimotor control of reaching. The outcomes of these three experiments can be summarized into 3 main points. First, the onset latency of the SLR does not change with increases in task complexity during any of the three experiments. Second, the SLR is directionally tuned to the stimulus location relative to the hand, not eye, position. Finally, the SLR magnitude is influenced by the pre-planned initial movement trajectory.

There are many similarities between the SLR’s visuomotor properties, which is evoked from a static posture, and rapid online corrective reaching movements to displaced visual (Gaveau *et al.*, 2014) or tactile stimuli (Pruszynski *et al.*, 2016). For example, the ∼100 ms latency of the SLR is consistent with previous reports of EMG response latencies to a displaced visual stimulus (Soechting & Lacquaniti, 1983; Fautrelle *et al.*, 2010), and occurs early enough to change reach kinematics within ∼150 ms (Carlton, 1981). Like the SLR, the latency of the online corrective movement is not modulated by changes in task demand (Oostwoud Wijdenes *et al.*, 2011; Franklin *et al.*, 2016). In an anti-reach paradigm, both the SLR (Gu *et al.*, 2016) and the initial trajectory of the corrective movements (Day & Lyon, 2000) are invariably directed towards the stimulus, even though the participants eventually moved in the opposite direction. Additionally, both the SLR (**Fig. 1**) and corrective movements (Diedrichsen *et al.*, 2004) are encoded in a hand-centric reference frame, reflecting stimulus location relative to the hand regardless of current eye position. Given the similarities between the SLR and corrective reach movements, we suggest that both are driven by a fast visuomotor system that lies in parallel to the well-studied corticospinal pathways (Alstermark & Isa, 2012).

It is tempting to speculate about the pathway that could be underlying the SLR, and by extension, corrective reach movements. Our findings are consistent with previous suggestions that corrective movements are mediated by visual inputs relayed through the superior colliculus (SC) via the reticulospinal pathway (Day & Brown, 2001; Reynolds & Day, 2012). For example, many neurons in intermediate and deep layers of the SC discharge a volley of action potentials with 50 ms of visual stimulus onset (Wurtz & Goldberg, 1972) that depends on the integrity of the lateral geniculate nucleus and primary visual cortex (Schiller *et al.*, 1979). Moreover, axons of these visually-responsive SC neurons contribute to the descending predorsal bundle that branches into the reticular formation (Rodgers *et al.*, 2006), leading to SLRs on neck muscles that promote orienting head movements (Corneil *et al.*, 2004, 2008; Rezvani & Corneil, 2008). In addition to its role in oculomotor control, the SC also plays a more general role in whole-body orienting (Gandhi & Katnani, 2011; Corneil & Munoz, 2014) and proximal limb control (Lünenburger *et al.*, 2001). Stimulation (Philipp & Hoffmann, 2014) and chemical inactivation (Song *et al.*, 2011) of the SC can influence reaching behavior in non-human primates, in line with human imaging studies of selective SC BOLD-activation during reaching tasks (Linzenbold & Himmelbach, 2012; Himmelbach *et al.*, 2013). Reach-related SC neurons can also exhibit similar short-latency visual responses (Song & McPeek, 2015), and movement-related activity correlates with recruitment of proximal limb muscle activity (Werner *et al.*, 1997; Stuphorn *et al.*, 1999). Furthermore, like the SLR, a subset of these neurons operate in a hand-centric reference frame (Stuphorn *et al.*, 2000).

Others have proposed that corrective movements are mediated through a cortical pathway, specifically via the posterior parietal cortex (PPC) (Desmurget *et al.*, 1999; Pisella *et al.*, 2000). The ∼100 ms latency of the SLR and its expression in hand-centric reference frame are both inconsistent with the known properties of PPC activity. For example, the SLR latency in the human limb occurs at, or around the same time, as the peak of the visual response of the monkey PPC (Snyder *et al.*, 1998). Most of these visual responses are also not encoded in a hand-centric reference frame that we observed with the SLR (Batista *et al.*, 1999; Buneo *et al.*, 2002). Thus, while the PPC may be involved in the later phases of online corrections (Franklin *et al.*, 2016), it seems unlikely that the PPC is involved in generating the SLR. Additionally, while primary motor cortex and premotor cortex do exhibit rapid visual transient responses (Kwan *et al.*, 1981; Weinrich & Wise, 1982), a recent study have suggested that these visual transient responses do not affect the neural output in both primary and premotor cortices (Stavisky *et al.*, 2017).

Finally, the results shown in Experiments 2 and 3 demonstrate that advanced planning of a movement trajectory can influence SLR magnitude. In both experiments, participants viewed an obstacle with which they either had to avoid or intersect for an extended period of time prior to the presentation of the visual stimulus. Moreover, the stimuli could only appear at a limited number of locations (2 and 3 for Experiments 2 and 3, respectively). The influence of such advanced planning on the SLR is particularly apparent in Catch Trials in Experiment 3, where ‘Congruent’ stimulus location in line with the initial phase of the planned curved trajectory evoked a larger SLR than ‘Incongruent’ stimulus location (**Fig. 4**). Previous neurophysiological studies have shown anticipatory build-up neural activity well before movement onset to both spatial and non-spatial cues throughout the primary (Tanji & Evarts, 1976; Confais *et al.*, 2012) and premotor cortices (Mauritz & Wise, 1986; Cisek & Kalaska, 2005), as well as within the PPC (MacKay & Crammond, 1987; Snyder *et al.*, 2006). These higher-order skeletomotor regions project directly to the SC (Fries, 1984, 1985; Distler & Hoffmann, 2015), providing a route by which planned movement trajectories could pre-set SC activity prior to stimulus onset.

## ACKNOWLEDGMENTS

We thank D. Park for assistance in data collection for Experiment 1 and H. Ohashi for programming assistance with the modified updated maximum likelihood procedure for Experiment 3. This work was supported by operating grants from the Natural Sciences and Engineering Research Council of Canada (NSERC) to BDC [RGPIN-311680], PLG [RGPIN-238338], J.A.P. [RGPIN-2015-06714], from the Canadian Institutes of Health Research to B.D.C. [MOP-93796], a NSERC Alexander Graham Bell Canada Graduate Doctoral Scholarship to C.G., and J.A.P. received a salary award from the Canada Research Chairs program.

